# Single cell analyses of cancer cells identified two regulatorily and functionally distinct categories in differentially expressed genes among tumor subclones

**DOI:** 10.1101/2023.04.22.537630

**Authors:** Wei Cao, Xuefei Wang, Kaiwen Luo, Yang Li, Jiahong Sun, Ruqing Fu, Qi Zhang, Ni Hong, Edwin Cheung, Wenfei Jin

**Author notes:** Correspondence (NH); (EC); (WJ).

## Abstract

To explore the feature of cancer cells and tumor subclones, we analyzed 101,065 single-cell transcriptomes from 12 colorectal cancer (CRC) patients and 92 single cell genomes from one of these patients. We found cancer cells, endothelial cells and stromal cells in tumor tissue expressed much more genes and had stronger cell-cell interactions than their counterparts in normal tissue. We identified copy number variations (CNVs) in each cancer cell and found correlation between gene copy number and expression level in cancer cells at single cell resolution. Analysis of tumor subclones inferred by CNVs showed accumulation of mutations in each tumor subclone along lineage trajectories. We found differentially expressed genes (DEGs) between tumor subclones had two populations: DEG_CNV_ and DEG_reg_. DEG_CNV_, showing high CNV-expression correlation and whose expression differences depend on the differences of CNV level, enriched in housekeeping genes and cell adhesion associated genes. DEG_reg_, showing low CNV-expression correlation and mainly in low CNV variation regions and regions without CNVs, enriched in cytokine signaling genes. Furthermore, cell-cell communication analyses showed that DEG_CNV_ tends to involve in cell-cell contact while DEG_reg_ tends to involve in secreted signaling, which further support that DEG_CNV_ and DEG_reg_ are two regulatorily and functionally distinct categories.

## Introduction

Single cell RNA-seq (scRNA-seq) is one of the most powerful technologies to study tumor microenvironment and profiles all cell types in tumor tissue, has significantly increased our knowledge about immune cell infiltration, tumor progression and facilitate the development of immunotherapy (Fan and Rudensky 2016; Li et al. 2017; Azizi et al. 2018; Zhang et al. 2018; Zhang et al. 2020; Qin et al. 2021; Zheng et al. 2021; Qi et al. 2022). Meanwhile, intratumoral heterogeneity is continuously posing a challenge to diagnosis and treatment of cancer, leading to drug resistance and disease recurrence (Bedard et al. 2013; Junttila and de Sauvage 2013; Dagogo-Jack and Shaw 2018). There are successful stories on distinguish the cancer cells from normal cells using gene expression matrix of scRNA-seq data (Tirosh et al. 2016; Qin et al. 2021), However, it is difficult to accurately infer tumor subclones using gene expression matrix of scRNA-seq data (Qin et al. 2021; Becker et al. 2022). On the other hand, single cell whole genome sequencing (scWGS) is particularly useful in inferring intratumoral heterogeneity and tumor subclones (Navin et al. 2011; Bedard et al. 2013; Kim et al. 2018). However, scWGS could not distinguish different cell types with normal genomes and is high cost, which hinders its wide application.

An alternative approach for inferring intratumoral heterogeneity is using copy number variations (CNVs) inferred from scRNA-seq data (Patel et al. 2014; Tirosh et al. 2016; Puram et al. 2017; Venteicher et al. 2017; Azizi et al. 2018; Maynard et al. 2020). E.g., Patel *et al*. inferred the CNVs using scRNA-seq data and identified intratumoral heterogeneity in glioblastoma (Patel et al. 2014). Puram *et al*. used the CNVs inferred from scRNA-seq to identify the intratumoral and intertumoral heterogeneity in head and neck cancer, based on which they found the partial epithelial-to-mesenchymal transition. (Puram et al. 2017). These success stories of identification of cancer cells and tumor subclones using scRNA-seq inferred-CNV is promising. However, most studies focused on how tumor subclones involved in complex tumor ecosystem (Richards et al. 2021; Wu et al. 2021a; Xiang et al. 2021; Xu et al. 2021). The features of cancer cells and tumor subclones is still not well investigation.

In this study, we characterized the features of cancer cells by comparing them with their epithelial cell counterparts using scRNA-seq data. We analyzed the relationship between gene copy number and expression level at single cell resolution. We used CNVs in each cell to infer the tumor subclones and their lineage trajectories. Most importantly, we found DEGs between tumor subclones had two populations, namely DEG_CNV_ and DEG_reg_. DEG_reg_ mainly locates in regions without CNVs and low CNV variation regions, and enriched in cytokine signaling genes. While DEG_CNV_ locates in high CNV variation regions and enriched in housekeeping genes and cell adhesion associated genes.

## Result

### Study design and cell atlas of human colorectal cancers

We designed a workflow to characterize the features of cancer cells and tumor subclones by integrating scRNA-seq and scWGS (**Figure 1A**). In brief, we generated scRNA-seq of tumor tissues, para-cancerous tissues and normal tissues from three colorectal cancer (CRC) patients diagnosed with proficient mismatch repair (pMMR) adenocarcinoma. We analyzed the scRNA-seq data of total 36 samples from 12 CRC patients after integrating data from Lee *et al*. (Lee et al. 2020). We further conducted scWGS on patient CRC#1, a patient with high quality scRNA-seq data. We finally integrated the scRNA-seq data and scWGS data to explore the feature of cancer cell and tumor subclones, thus potentially provide biological insight into the gene regulation in cancer.

**Figure 1.**
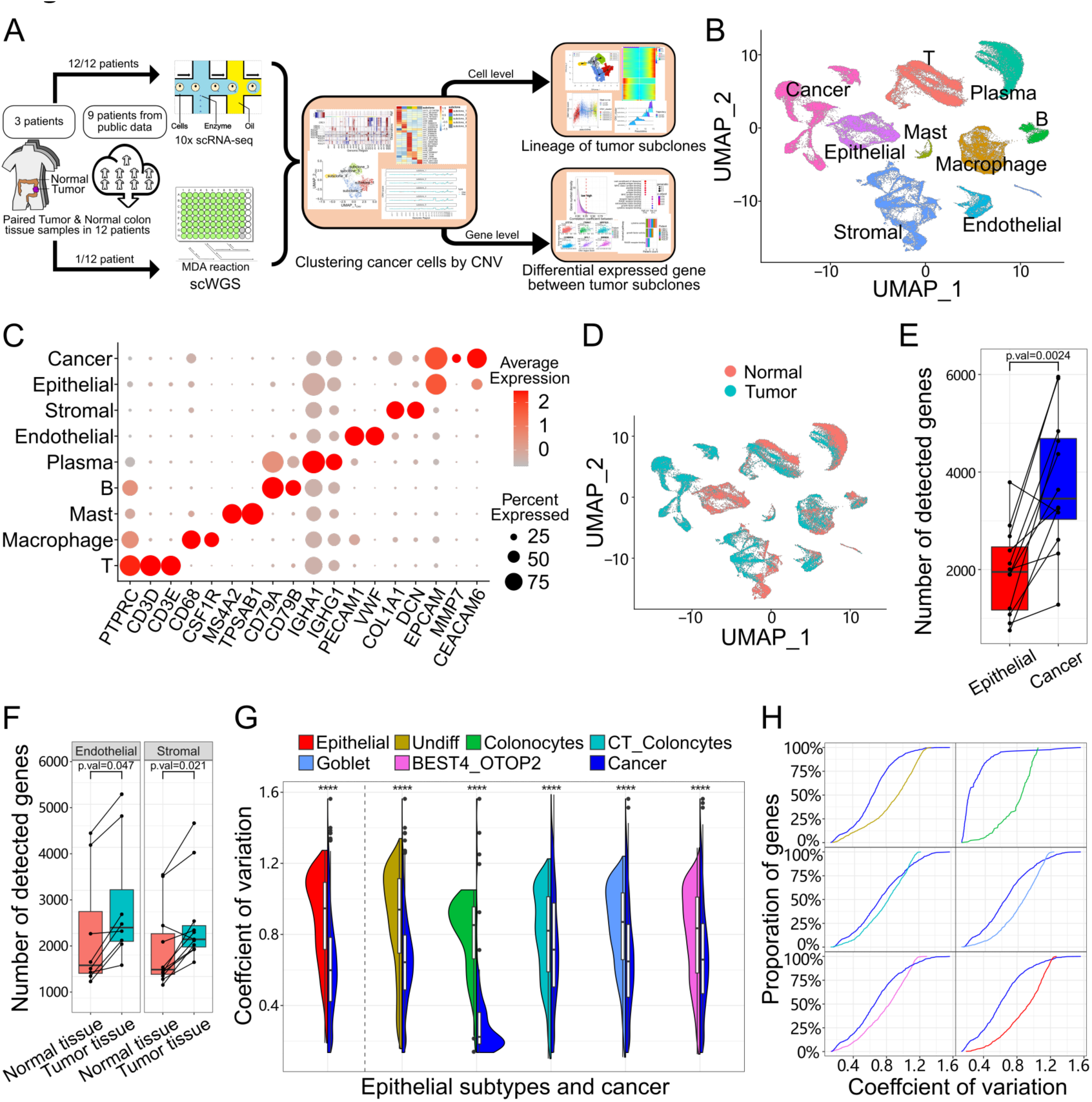
Cell atlas of CRC and increased cell activities in tumor tissues. **(A)** The scheme of this study. **(B)** UMAP visualization of 101,065 cells from tumor tissue and normal colorectal tissue in 12 patients, colored by cell type. **(C)** Dot plot of normalized expression level and expression percentage of cell-type-specific genes in 9 cell types. **(D)** UMAP visualization of 101,065 cells from tumor tissue and normal colorectal tissue, colored by tissue. **(E)** Box plot of average number of detected genes in each patient for cancer cells and epithelial cells. Wilcoxon signed rank test were performed to examine whether the number of detected genes is significantly different between cancer cells and epithelial cells. **(F)** Box plot of average number of detected genes in each patient in normal tissue and tumor tissue for endothelial cells and stromal cells. Wilcoxon signed rank test were performed to examine whether the number of detected genes were significantly different between normal tissue and tumor tissue for endothelial cells and stromal cells. **(G)** Splited violin plot of coefficient of variation (CV) of gene expression level of detectable genes among cells in epithelial cells and epithelial subtypes compared with the same genes in cancer cells in patients. Paired Wilcoxon signed rank test was used to examine whether the differences were significant. **(H)** Accumulation curve of CV of gene expression level in epithelial cells and epithelial subtypes compared with same genes in cancer cells in all patients.

After quality control, we obtained a total of 101,065 cells, with an average of 16,085 unique molecular identifiers (UMIs) and an average of 2,469 detected genes per cell (**Figure S1A**). Cells from para-cancerous tissues almost completely overlap with cells from normal tissues in UMAP plot, as well as cells from tumor border tissue almost completely overlap with cells from tumor tissue (**Figure S1B**), indicating these cells from different samples are similar to each other for each patient and batch effects among these samples could be ignored. Therefore, we merged cells in para-cancerous tissues to cells in normal tissues in each patient, and merged cells in tumor border tissues to cells in tumor tissues in each patient as well. The 101,065 filtered cells were clustered into nine cell types, namely cancer cells, T cells, macrophages, mast cells, B cells, plasma cells, endothelial cells, stromal cells and epithelial cells (**Figure 1B and C; Figure S1C**). For each cell type except cancer cells, cells in tumor tissues and their counterparts in normal tissue partially overlap on UMAP. Cancer cells did not overlap with their normal counterparts (epithelial cells) on UMAP plot, indicating the differences between them are pronounced (**Figure 1D**).

## Comparison of cells from tumor tissues with their counterparts in normal tissues

We found cancer cells had much more detected UMIs and more detected genes than their epithelial cell counterparts (**Figure 1E; Figure S1D**), potentially indicating cancer cells have increased cell size and expressed more genes after epithelial cells cancerization. Endothelial cells and stromal cells in tumor tissues also have significantly more detected UMIs and more detected genes than their counterparts in normal tissues (**Figure 1F; Figure S1E**), which may be caused by the different microenvironment between normal tissues and tumor tissues. While detected UMIs and detected genes of other cell types in tumor tissues were not significantly different from their counterparts in normal tissues (**Figure S1F and G**).

We identified total 600 DEGs between cancer cells and epithelial cells, of which 357 genes were cancer cell specific and 243 genes were epithelial cell specific (**Figure S2A**). The cancer cells specific genes were enriched in ribonucleoprotein complex biogenesis, protein folding, neutrophil degranulation, regulation of apoptotic signaling pathway, cytokine signaling, VEGF pathway and MYC pathway (**Figure S2A and B**), potentially indicating cancer cells have increased cell activity, and promotion of immune cell infiltration/maturation. The epithelial cell specific genes were significantly enriched in response to bacterium, metallothioneins bind metals, transport of small molecules, epithelial cell differentiation, response to hormone, primary alcohol metabolic process, regulation of defense response and response to nutrient (**Figure S2A and C**), potentially indicating epithelial cells are more sensitive to external environments than their cancer cell counterparts.

We identified total 204 DEGs between tumor associated endothelial cells and normal endothelial cells, of which 100 genes were tumor associated endothelial cell specific and 104 genes were normal endothelial cell specific (**Figure S3A**). The tumor associated endothelial cell specific genes were enriched in extracellular matrix organization, blood vessel development, regulation of angiogenesis and regulation of cell-substrate adhesion (**Figure S3A and B**), indicating endothelial cells in tumor played important role in promotion of angiogenesis. The normal endothelial cell specific genes were enriched in negative regulation of cell adhesion, positive regulation of cell migration, negative regulation of cell population proliferation and negative regulation of catalytic activity (**Figure S3A and C**), indicating normal endothelial cell have weaker cell adhesion and proliferation than their counterparts in tumor tissue. We further identified total 310 DEGs between tumor associated stromal cells and normal stromal cells, of which 164 genes were tumor associated stromal cell specific and 146 genes were normal stromal cell specific (**Figure S3D**). The tumor associated stromal cell specific genes were enriched in extracellular matrix organization, vasculature development and response to wounding (**Figure S3D and E**), while normal stromal cell specific genes were enriched in negative regulation of cell population proliferation, negative regulation of cell adhesion and response to hypoxia (**Figure S3D and F**).

### Cancer cells have lower cell-cell variation than their epithelial cells

Although the high cellular heterogeneity of tumor tissue is well known, we found the cell-cell variations of cancer cells are significantly lower than that of epithelial cells (**Figure 1G**). To further explored cell-cell variation, we further cluster epithelial cells into several subsets (**Figure S4A and B**) using previously reported marker genes (Parikh et al. 2019). The cell-cell variations of cancer cells are significantly lower than that of each epithelial cell subset in pooled cells or in each CRC patient (**Figure 1G; Figure S4C**). The low cell-cell variation of cancer cells may be due to cancer cells were derived from the clone expansion of one or a few mutated cells, thus the cancer cells had a few states. To understand the cell-cell variation at gene level, we compared the cell-cell variation of each gene between epithelial cells and cancer cells. We found cell-cell variations of most genes in epithelial cells were much higher than that in cancer cells (**Figure 1H; Figure S4D**), consist with aforementioned observations that cell-cell variations of epithelial cells are higher than that of cancer cells.

### Heterogeneity of cancer cells and inference of CNVs

To further understand the feature of cancer cells, we conducted UMAP analysis on epithelial-like cells including cancer cells and epithelial cells. We found epithelial cells from normal tissue in all patient samples mixed in a single cluster, while the cancer cells from each patient have a patient-specific cluster (**Figure 2A**). The results indicate cancer cells are quite different from patient to patient, implying high interpatient diversity or interpatient heterogeneity. Although the high inter-patient heterogeneity of cancer cells, the entropy of cancer cells from each patient is significantly higher than their epithelial cell counterparts (**Figure S5A**), potentially indicating de-differentiation or increased stemness of cancer cells after epithelial cell carcinogenesis. Indeed, the cancer cells have much higher stemness scores than their epithelial cell counterparts (**Figure S5B**). Since the inter-patient differences among cancer cells are dominant, we inferred copy number variations (CNVs) in each patient using scRNA-seq data. In order to better present CNVs, we separated the genomic regions into duplication regions, deletion regions and regions without CNVs (noCNV) in each patient according to CNV level of the genomic regions in cancer cells (**Figure 2B**).

**Figure 2.**
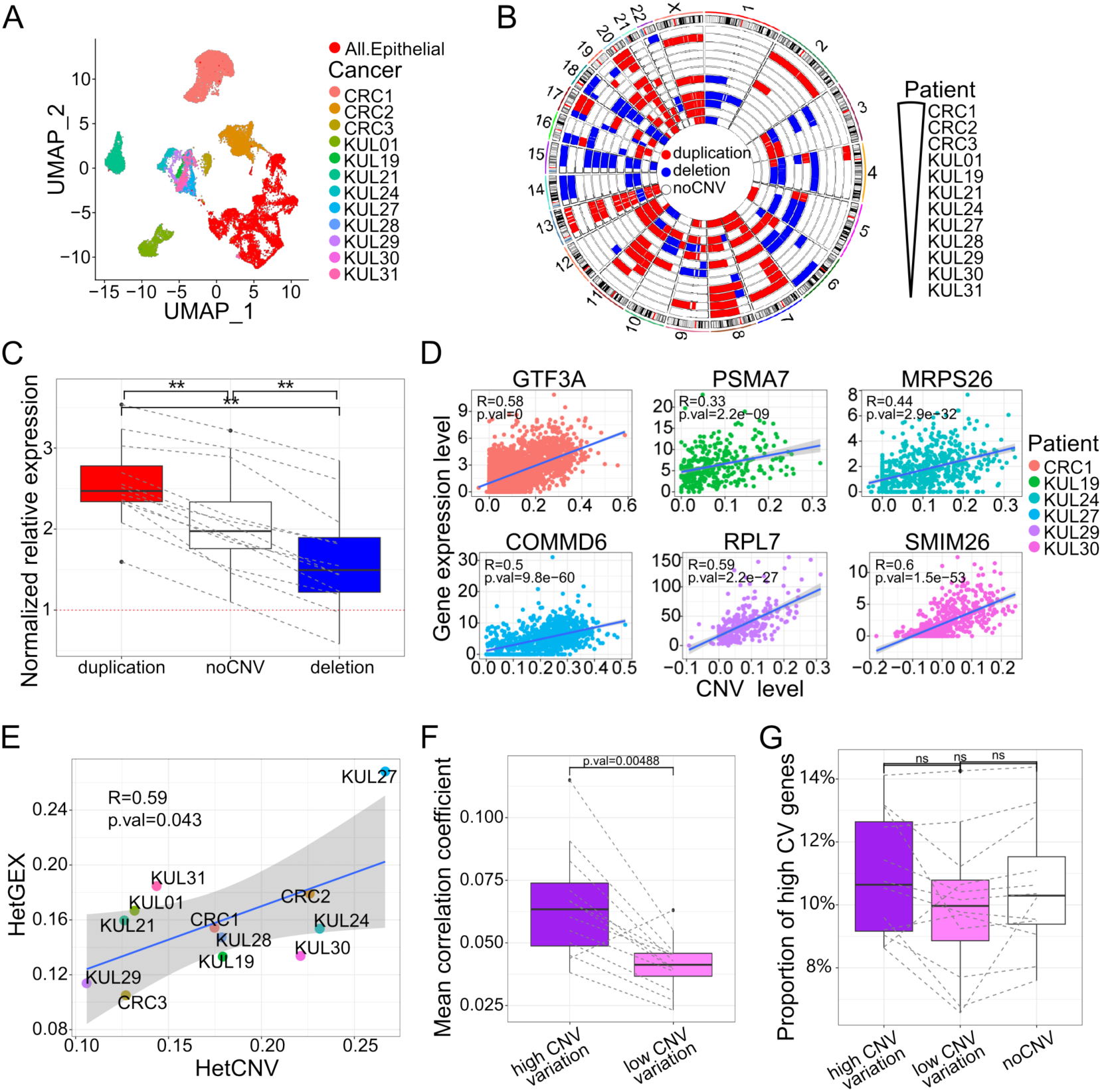
Cancer cells have a specific regulatory mechanism that gene expression levels depend on its DNA copy number. **(A)** UMAP visualization of epithelial cells (red) and cancer cells, with cancer cells colored by patient. **(B)** Circos heatmap showing duplication, deletion, and noCNV genomic regions in 12 patients. **(C)** Box plot of gene expression in cancer cells normalized to their expression in normal counterpart in duplication, deletion, and noCNV genomic regions in each patient. Paired Wilcoxon signed rank test was used to examine whether the differences were significant. **(D)** Scatter plots showed the correlation between expression level and CNV level in CNV regions. Shaded areas corresponded to 0.95 confidence interval. **(E)** Scatter plot of the correlation between Intratumoral heterogeneity scores based on gene expressions (HetGEX) and Intratumoral heterogeneity scores based on CNVs (HetCNV) for the 12 patients. Shaded areas corresponded to 0.95 confidence interval. **(F)** Box plot of correlation coefficient between gene expression level and CNV level in CNV variation regions and low CNV variation regions. Paired Wilcoxon signed rank test was used to examine whether the differences were significant. **(G)** Box plot of the proportion of high CV genes between chromosome regions. chromosome regions were classified into high CNV variation, low CNV variation region and no CNV region in each patient. Paired Wilcoxon signed rank test was used for examining whether differences were significant between each chromosome region pairs.

### Gene expression level correlated with its DNA copy number in cancer cells

The expression level of each gene in cancer cells was normalized using its expression level in epithelial cell counterparts. The genes located in duplication regions express significantly higher than the genes in noCNV regions (P=4.9×10^-4^) and deletion regions (P=4.9×10^-4^), with genes in deletion regions have the lowest expression levels (**Figure 2C**). These results indicate that gene expression in cancer cells are strongly associated with its DNA copy number. Indeed, there are many genes showing strong correlation between expression level and CNV level at single cell resolution (**Figure 2D**). We further calculated intra-tumoral heterogeneity scores based on CNV level (HetCNV) and intra-tumoral heterogeneity scores based on gene expression (HetGEX). We found HetGEX highly correlated with HetCNV (**Figure 2E**), potentially indicating the heterogeneity of CNV impacts the heterogeneity of gene expression. The copy number of CNV and heterogeneity of CNV influenced the gene expression should be a unique features of cancer cells since normal cells are diploid and gene expression were precisely regulated.

### Gene expression levels are more likely to be determined by its DNA copy number in high CNV variation regions

To further investigate the relationship between gene expression level and CNV level, CNV regions were classified into high CNV variation regions and low CNV variation regions in each patient. We found genes in high CNV variation regions have significant higher correlation coefficient between gene expression level and CNV level than low CNV variation regions (**Figure 2F**). Meanwhile, the fractions of genes showing high expression variation in duplication regions, noCNV regions and deletion regions are not significantly different (**Figure 2G**), indicating genes with high expression variation have no bias in different CNV region categories.

### Inference of tumor subclones using inferred CNVs

In order to better understand the heterogeneity of cancer cells, we inferred copy number variations (CNVs) in each epithelial-like cell using scRNA-seq data. Indeed, the epithelial cells from normal tissues did not have pronounced the chromosomal deletions or duplications; while the cancer cells from different patients have many patient-specific genomic alterations over large genomic regions (**Figure 3A**), which could explain why cancer cells showed patient specific cluster on UMAP plot. Although the differences of CNVs among patients are major, we also found the differences of CNVs among cancer cells from the same patient (**Figure 3A**). To better illustrate the heterogeneity of cancer cells in each patient, the inferred CNVs was used to cluster cancer cells into tumor subclones in each patient (**Figure S5C**).

**Figure 3.**
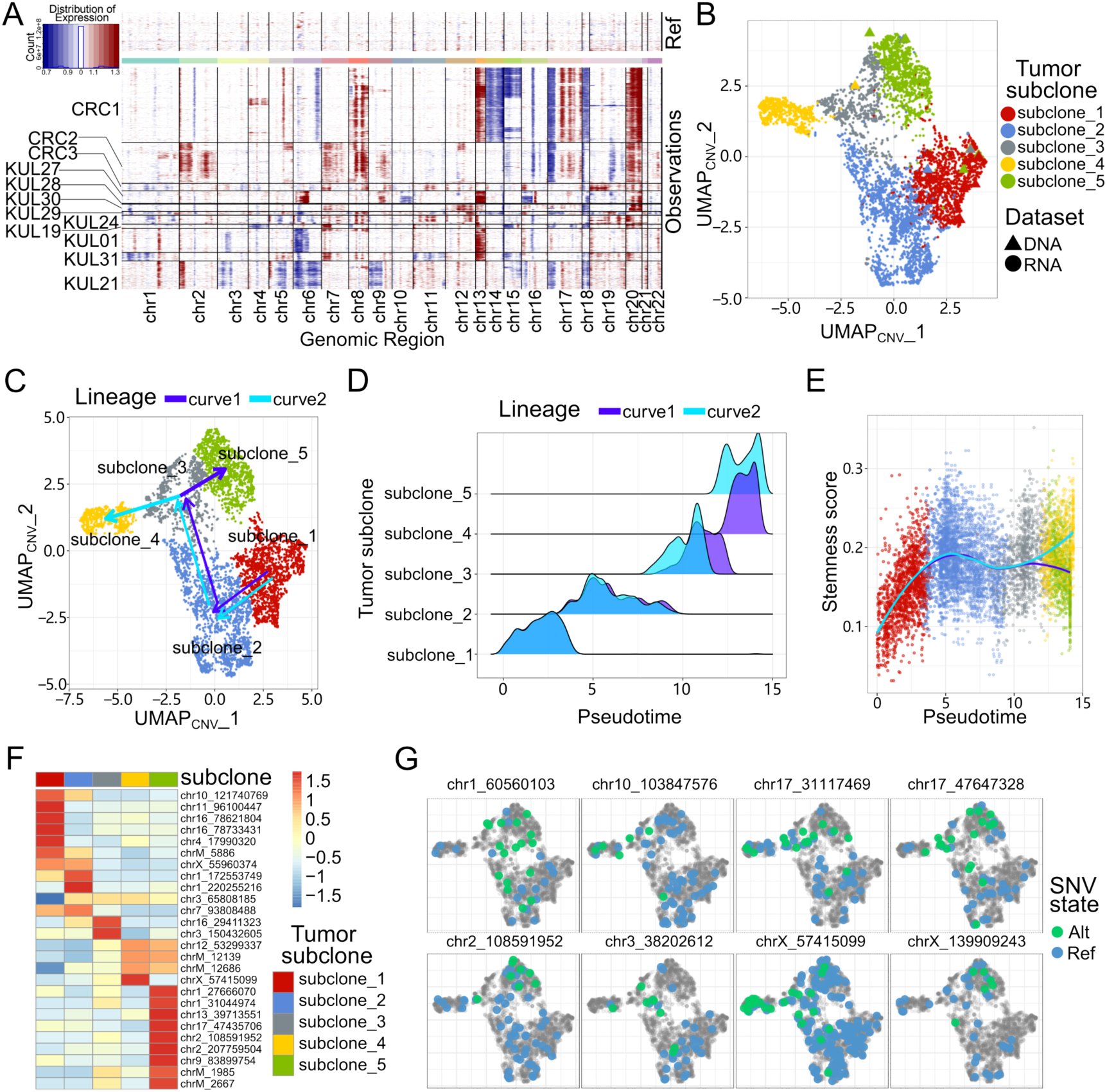
Tumor subclones and its pseudotime trajectory revealed by single cell CNV analyses. **(A)** Heatmap of CNV level in reference normal cells and cancer cells, with rows and columns representing cells and genomic loci, respectively. **(B)** UMAP visualization of cancer cells from both scWGS and scRNA data in patient CRC#1, Cells colored by tumor subclone inferred by CNVs and shaped by datasets. **(C)** Lineage trajectories of cancer cells in CRC#1 on UMAP. **(D)** Density plot of the positioning of cells from each tumor subclone on pseudotime trajectories. **(E)** Scatter plot of the stemness score along pseudotime trajectories showed the stemness increase quickly at the early stage. **(F)** Heatmap of SNV alternative ratios’ z-score by row in each tumor subclone. **(G)** UMAP visualization of cancer cells in CRC#1, colored by the SNV alternative ratio of each SNV.

We also inferred CNVs and tumor subclones using scWGS data of patient CRC#1. We found that the CNVs inferred by scRNA-seq in each tumor subclones were essentially consistent with that inferred by scWGS (**Figure S6A-C**), although CNVs inferred in scWGS were more accurate and higher resolution than that inferred by scRNA-seq. Furthermore, cell abundance of non-cancer cells and each tumor subclone inferred based on scRNA-seq data is very similar to its tumor subclone counterpart inferred by scWGS data (**Figure 3B; Figure S6D**).

### Tumor lineages and accumulation of mutations in each tumor subclone

We inferred pseudotime of cancer cells to better understand the relationship among these tumor subclones. We analyzed the CNV data using slingshot and identified two lineage trajectories of tumor subclones: (1) subclone 1 → subclone 2 → subclone 3 → subclone 4; (2) subclone 1 → subclone 2 → subclone 3 → subclone 5 (**Figure 3C and D**). Furthermore, the two main lineages inferred by monocle2 were essentially consistent with that inferred by slingshot (**Figure S6F-H**). For the inferred lineages, the stemness scores of cancer cells increase rapidly at the beginning of the pseudotime. After separation of the two lineages, the scores continuously increase along lineage #2 while scores along lineage #1 are stable (**Figure 3E**). Furthermore, the inferred CNV scores of each tumor subclone increased along with lineage trajectories (**Figure S6E**), indicating the accumulation of CNV along lineages.

We further called single nucleotide variants (SNVs) in scRNA-seq data using GATK. We clustered alternative ratio of SNVs in tumor subclones and identify many tumor subclones specific SNVs (**Figure 3F**). The distribution of alternative alleles on UMAP showed tumor subclones specific (**Figure 3G**), which support the accumulation of mutations in each tumor subclone.

### The DEGs between tumor subclones have two populations

We identified 288 DEGs and 85 DEGs between tumor subclones by setting average expression level fold change >1.5 and >1.8, respectively (**Figure 4A**). The DEGs are significantly enriched in interferon signaling, neutrophil degranulation, cellular response to cytokine stimulus and regulation of cell-cell adhesion (**Figure S7A and B**). The genes showing the most significant difference include *PLCG2* and *CXCL14*, which have been reported as tumor-associated marker genes (Sjoberg et al. 2016; Li et al. 2021). The distribution of correlation coefficient between gene expression level and CNV level revealed the existence of two populations of DEGs: DEGs with high CNV-expression correlation and DEGs with low CNV-expression correlation (**Figure 4B**). The natural separation between the two populations of DEGs may indicate they have different features.

**Figure 4.**
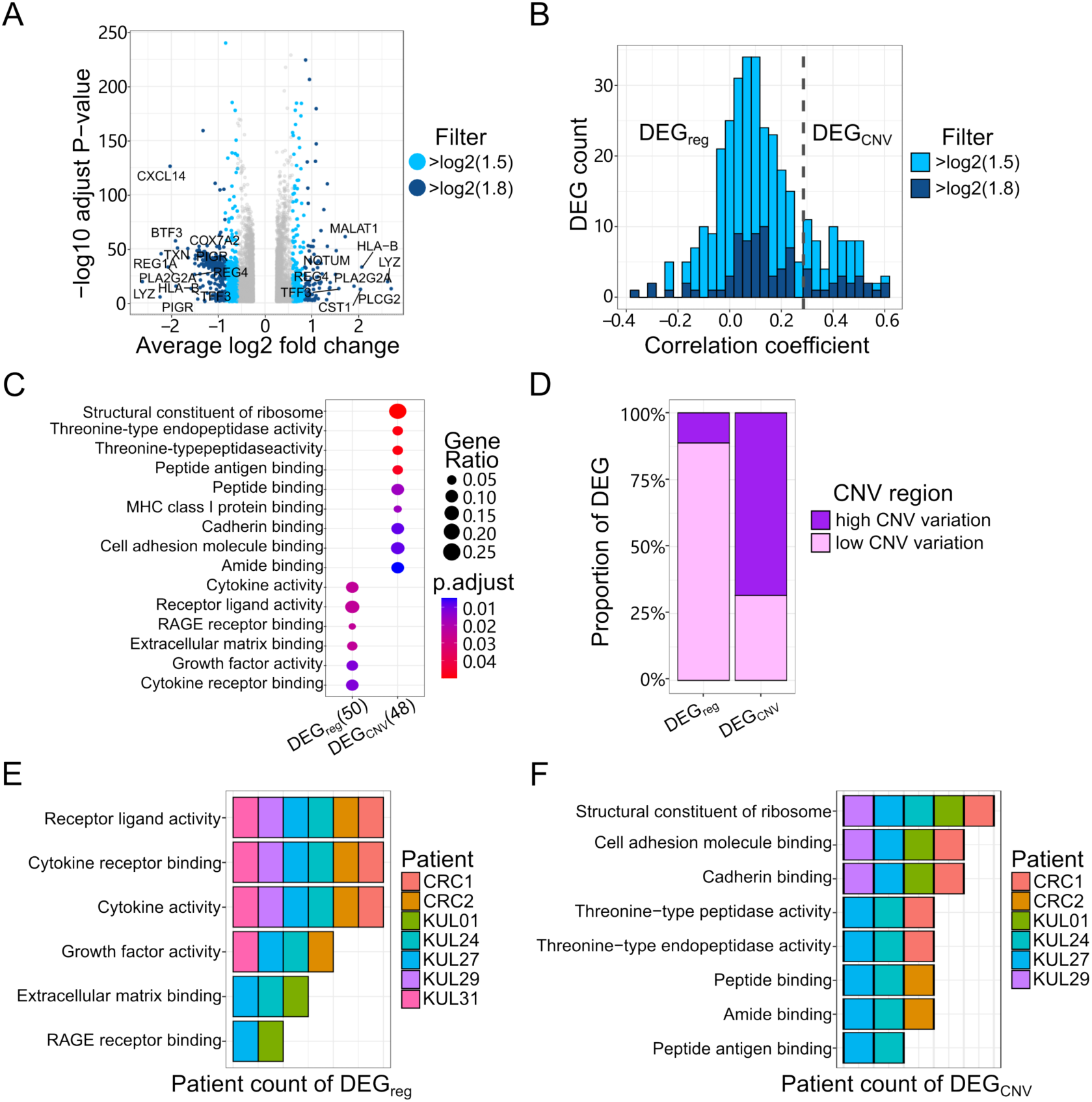
DEGs between tumor subclones showed two regulatorily and functionally distinct populations. **(A)** Volcano plot of DEGs between tumor subclones in each patient. Light blue and blue represent fold change > 1.5 and fold change > 1.8, respectively. **(B)** Histogram plot of correlation coefficient between gene expression level and CNV level. The DEGs sets were set at fold change >1.5 and fold change >1.8, respectively. **(C)** GO enrichment analysis of DEGs with high CNV-expression correlation (DEG_CNV_) and DEGs with low CNV-expression correlation (DEG_reg_). Dot size and color represents gene ratio and adjusted p-value, respectively. **(D)** Percentage of genes located in high CNV variation region and low CNV variation regions in DEG_CNV_ and DEG_reg_. Purple and pink represents high CNV variation regions and low CNV variation regions. **(E)** Bar plot of patient count representing whether the patient’s DEG_reg_ is enriched in the GO term. **(F)** Bar plot of patient count representing whether the patient’s DEG_CNV_ is enriched in the GO term.

### DEGs with high or low CNV-expression correlation enrich different GO terms

GO analysis showed DEGs with high CNV-expression correlation were enriched in housekeeping genes and receptor-ligand binding genes such as peptide antigen binding, (P=3.0×10^-3^), cadherin binding (P=0.041), structural constituent of ribosome (P=2.6×10^-11^) and MHC class I protein binding (P=0.033) (**Figure 4C**), which indicate differential expression of these housekeeping genes between tumor subclones depend on the change of their DNA copy number. While DEGs with low CNV-expression correlation were enriched in secretory signaling such as cytokine activity (P=0.026), RAGE receptor binding (P=0.026), cytokine receptor binding (P=0.038) and growth factor activity (P=0.038) (**Figure 4C**), indicating differential expression of these cytokine signaling genes do not depend on the change of their DNA copy number.

Indeed, DEGs with high CNV-expression correlation are mainly in high CNV variation regions (**Figure 4D**), whose expression level are strongly depended on CNV level thus called DEG_CNV_. While DEGs with low CNV-expression correlation are mainly in noCNV regions and low CNV variation regions (**Figure 4D**), which may be caused by differences of gene regulation thus called DEG_reg_. Ribosome genes, belonging to housekeeping genes, were pronounced enriched in DEG_CNV_ comparing with that in DEG_reg_ (**Figure S7C**). Furthermore, the genes in DEG_CNV_ were expressed higher and were detected in much higher fraction of cells (**Figure S7D-F**), which is consistent with previous report that housekeeping genes expressed much higher than other genes (Jin et al. 2012).

Analyses of signaling enriched in DEG_reg_ showed that cytokine activity was detected in most patients (**Figure 4E**), indicating the change of expression of cytokine signaling genes, without gene copy change, played important roles in shaping tumor subclones or tumor heterogeneity in many patients. On the other hand, analyses of signaling enriched in DEG_CNV_ showed that structural constituent of ribosome, cell adhesion molecule binding and cadherin binding in several patients (**Figure 4F**), indicating ribosome and cell adhesion molecule binding genes played important roles in shaping tumor subclones or tumor heterogeneity in many patients. The differences of molecule binding and the cytokine signal pathways between tumor subclones should be two quite different mechanisms for shaping tumor subclones differences.

### DEG_CNV_ and DEG_reg_ have different cell-cell communication pattern

Cell-cell communications between most cell types in tumor tissue are significantly higher than that in normal tissue (**Figure 5A and B; Figure S8A**). Among all cell-cell communication pairs, the 3 top increased communication pairs are endothelial-cancer/epithelial, stromal-cancer/epithelial, endothelial-stromal (**Figure 5B**), suggesting cell-cell communication among the non-immune cell pairs increased the most. Further analyses showed that cell-cell communications between immune cell and non-immune cell increased the second highest, while cell-cell communications between immune cell and immune cell increased but the least (**Figure 5B; Figure S8B**). We further compared the cell-cell communication between DEG_CNV_ and DEG_reg_. The cell-cell communication of DEG_CNV_ and DEG_reg_ are mainly cell-cell contact and secreted signaling, respectively (**Figure 5C and D**), which is consistent with our aforementioned enrichment analysis that DEG_CNV_ and DEG_reg_ enriches in cell adhesion and cytokine signaling genes, respectively. We found both the cell-cell interaction weight of cytokine signaling and cell-cell interaction weight of adhesion signaling in tumor tissue are significantly higher than that in normal tissue (**Figure 5E**). There are total 11 pathways significant changed between tumor tissue and normal tissue. Among those pathways, six of them shared between cytokine signaling and adhesion signaling, four of which are significant increased (FN1, VISFATIN, THBS and TGFb) and two of which are significant decreased in tumor tissue (PTN and CXCL) (**Figure 5F**). Other pathways are all significantly increased in tumor tissue. MIF is the only cytokine signaling specific pathway that significant increase the interaction between immune cell with cancer/epithelial cell. Among the adhesion signaling specific pathway, COLLAGEN, LAMININ, MK and CDH1 among most cell type pairs are significantly higher in tumor tissue than in normal tissue (**Figure 5F; Figure S8C**). Furthermore, CDH1 is mainly released and received by cancer/epithelial cells, MIF is mainly received by macrophage and T cells, and COLLAGEN, LAMININ, MK mainly released by stromal and endothelial cell (**Figure 5G-I; Figure S8D and E**), which potentially indicate DEG_reg_ mainly affect immune system and DEG_CNV_ mainly communicate with microenvironment and cancer cells themselves.

**Figure 5.**
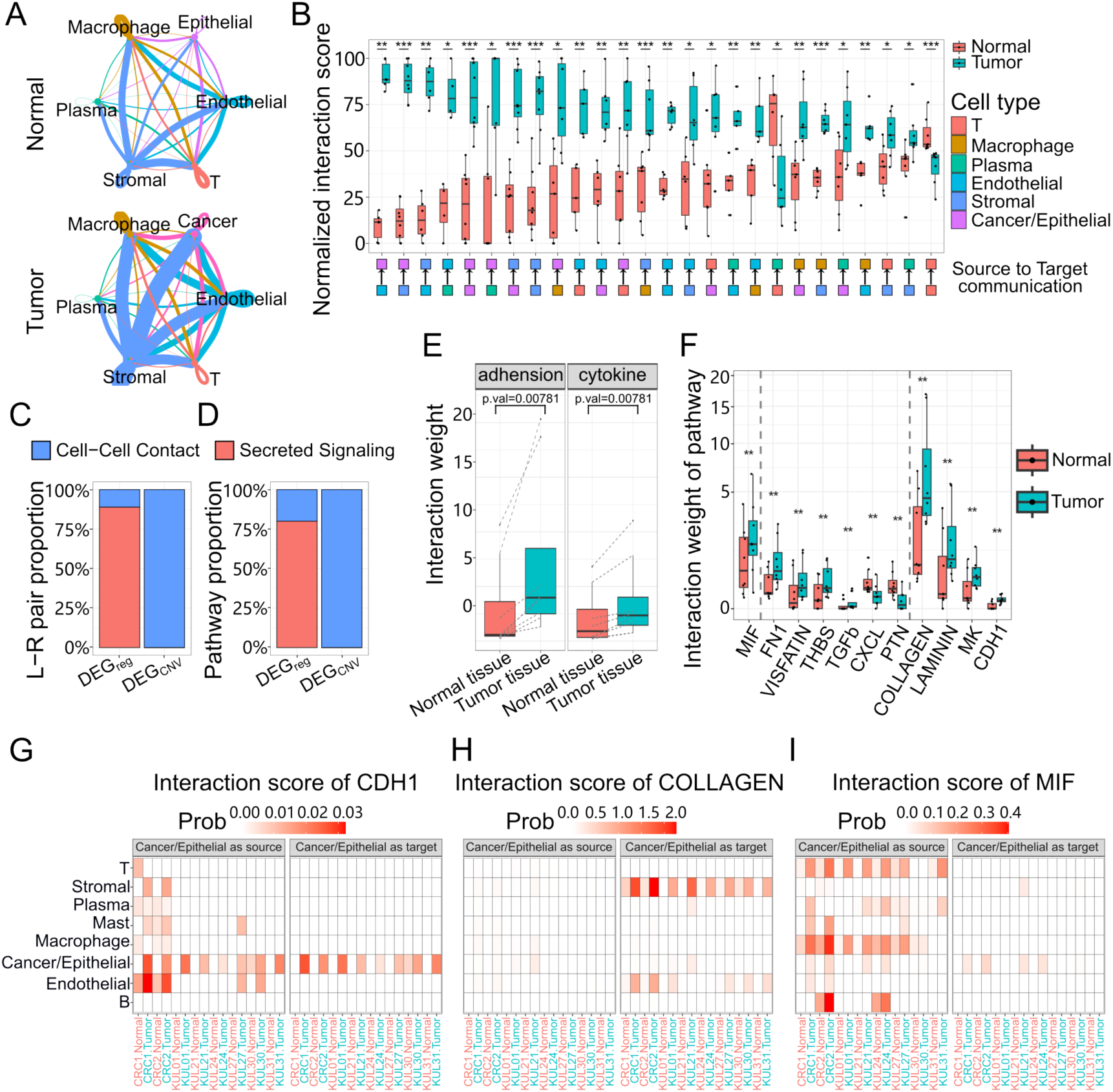
The change of cell-cell communication between tumor tissues and normal tissues. **(A)** Cell-cell communication networks of ligand-receptor pairs in normal tissue (upper panel) and tumor tissue (lower panel) in all patients. Edge colors are consistent with the signaling senders. Edge weights are proportional to the interaction weight, and thicker edge line indicates a stronger signal. **(B)** Box plot of normalized interaction score of cell type pairs in all patients. Pairs are ranked by the interaction difference between normal tissue and tumor tissue. Paired Wilcoxon signed rank test was used to examine whether the differences between tumor tissue and normal tissue were significant in each source to target communication pairs, p values were adjusted by BH. **(C)** Bar plot of percentage of genes involved in cell-cell contact and secreted signaling in DEG_CNV_ and DEG_reg_. **(D)** Bar plot of percentage of pathways involved in cell-cell contact and secreted signaling in DEG_CNV_ and DEG_reg_. **(E)** Box plot of interaction weight of cytokine and adhesion signaling pairs in tumor tissue and normal tissue. Paired Wilcoxon signed rank test was used for examining whether differences were significant. **(F)** Box plot of interaction weight of 6 cytokine and adhesion shared pathways (FN1, VISFATIN, THBS and TGFb; PTN and CXCL), one cytokine specific pathway (MIF), four adhesion specific pathways (COLLAGEN, LAMININ, MK and CDH1) that in tumor and normal tissue. Wilcoxon signed rank test was used for examining whether differences were significant between tumor and normal tissue in each pathway, p values were adjusted by BH. **(G-I)** Heatmap of interaction score of cancer/epithelial cell to other cell types (left) and other cell types to cancer/epithelial cell (right) of CDH1 **(G)**, COLLAGEN **(H)** and MIF **(I)** pathways in each patient’s normal and tumor tissue.

## Discussion

It is well known that cancer cells are quite different from their normal counterparts, but the *in vivo* differences between cancer cells and their normal counterparts have not been systematically studied at single cell resolution. In this study, we found the cancer cells expressed much more genes than their normal epithelial cell counterparts. We found cancer cells highly expressed genes associated with protein folding, neutrophil degranulation, cellular responses to stress, cytokine signaling, VEGF pathway and MYC pathway, indicating cancer cells have increased cell activity and enhanced of immune cell infiltration/maturation compared with their normal counterpart. The results also indicated that cancer cells expressed much more genes, and stronger cell-cell communication with other cell types than their normal counterparts. Furthermore, the stromal cells and endothelial cells in tumor tissue also expressed much more genes and had much stronger cell-cell communication with other cell types than their counterparts in normal tissue. Stromal cells and endothelial cells in tumor tissue highly expressed genes associated with extracellular matrix organization, response to wounding and angiogenesis. Therefore, cancer cells and stromal cells played quite different role in shaping the tumor microenvironment.

The dramatic differences between the cells in tumor tissue and their counterpart in normal tissue could be attributed to their different tissue microenvironments. Interestingly, we found the strong correlation between gene expression and its DNA copy number in cancer cells, indicating gene expression in cancer cells are more likely to depend on its DNA copy number. This observation is consistent with the recent studies reported the highly expressed genes in tumor usually have a lot DNA copies in the form of extrachromosomal circular DNA (eccDNA) (Turner et al. 2017; Wu et al. 2019). Considering that most genes have two copies in mammalian cells while their expression levels vary greatly, thus the copy number of DNA should not be a strong factor affecting the difference of gene expression in normal cells. In short, the CNV-expression correlation should be a very specific and unique features of gene regulation in cancer cells.

Cancer cells usually were classified into different cell subsets based on gene expression matrix of scRNA-seq (Li et al. 2017; Chowdhury et al. 2021; Sacchetti et al. 2021; Sun et al. 2021; Wang et al. 2021; Xiang et al. 2021; Zowada et al. 2021). In this study, we showed CNVs inferred by scRNA-seq data were essentially consistent with that inferred by scWGS data. We inferred the tumor subclones using the CNVs inferred by scRNA-seq and found these tumor subclones have subclone specific SNVs, which further support the reliability of the inferred tumor subclones. We further identified the tumor subclones and inferred the lineage relationship of these tumor subclones in CRC#1, which support the accumulation of mutations in each tumor subclone. We further identified the DEGs among tumor subclones and found these DEGs had two populations according to CNV-expression correlation. Interestingly, DEG_CNV_, mainly located in high CNV variation regions, has high CNV-expression correlation and enriches in housekeeping genes and adhesion signaling genes. While DEGreg, mainly located in noCNV regions and low CNV variation regions, has low CNV-expression correlation and enriched in secretory signaling particularly cytokine signaling, which changed cell communication between cancer cell with immune cells. These results indicate that cytokine signaling genes showed tumor subclones difference but these differences were not determined by its DNA copy but by gene regulation. While housekeeping genes and adhesion associated genes that showed tumor subclones difference between tumor subclones are more likely to depend on change at its DNA level.

In conclusion, we characterized the features of cancer cells in tumor tissues by comparing with their counterparts in normal tissues at single cell resolution. We found that cancer cells expressed much more genes, had stronger metabolic activities and stronger cell-cell interaction with other cell types than their normal counterparts. In particular, we found the strong correlation between gene expression and its DNA copy number in cancer cells, potential indicating CNV-expression correlation is a very specific and unique features of gene regulation in cancer cells. We identified the tumor subclones using CNVs and inferred the lineage relationship of these tumor subclones in CRC1, which support the accumulation of mutations in each tumor subclone. We found two DEGs populations among tumor subclones according to CNV-expression correlation, DEG_CNV_ and DEGreg. DEGreg has low CNV-expression correlation and enriched in cytokine signaling genes, while DEG_CNV_ has high CNV-expression correlation enriched in housekeeping genes and adhesion signaling genes.

## Materials and Methods

### Collection of colorectal cancer samples

This study was approved by IRBs at Southern University of Science and Technology (SUSTech). Three patients diagnosed with colorectal cancer were enrolled in the First Affiliated Hospital, SUSTech. All patients signed informed consent form approved by the IRB of SUSTech. None of them received chemotherapy, radiation or immunotherapy prior to tumor resection.

### Tissue dissociation and cell suspension

Tissue obtained immediately after tumor resection was used to prepare cell suspension. In brief, freshly resected tissue was washed with RPMI 1640 (Thermo Fisher Scientific) and minced with scissors. The tissues were digested with Human Tumor Dissociation Kit (Miltenyi Biotec) according to manufacturer’s manual. Dissociated cells were passed through a 70μm cell sieve and dead cells were removed using Dead Cell Removal Kit (Miltenyi, 130-090-101). After centrifugation with 300g at 4°C for 5 minutes, the supernatant was discarded and the cells were washed with PBS. Then, 1ml cold PBS containing 2% FBS was added into the tube to suspend the cells. The cell suspension was used for further analysis.

### scRNA-seq

scRNA-seq libraries were prepared using Chromium Single Cell 3ʹ v3 Reagent Kits (10x Genomics) following manufacturer’s protocols. In brief, approximately 1.6×10^4^ cells were loaded into each lane to produce the Gel Beads-in-Emulsions (GEMs), followed by reverse transcription, cDNA amplification and library construction. Qubit and Qsep100 were used to measure DNA concentration and fragment length of each library, respectively. The scRNA-seq libraries were sequenced on Illumina NovaSeq6000, instrument with pair-end sequencing and dual indexing according to recommended Chromium platform protocol.

### Single-cell DNA sequencing (scWGS) and bulk DNA-seq

scWGS library were prepared using QIAseq FX Single Cell DNA Library Kit. Cell suspension of tumor tissue and normal tissue from patient CRC#1 were used for library preparation. In brief, 4ul PBS was added to each well of a 96-well plate. Single cells were sorted into individual well of the 96-well plate containing PBS and 3μl Buffer D2. Cells were incubated for 10 min at 65°C and the reaction was stopped by adding 3 μl of Stop Solution. For each amplification reaction, 40 μl master mix was added to the 10 μl reaction to denature the DNA, followed by incubation at 30°C for 2 h and stopping by incubating at 65°C for 3 min. In this way, we completed genomic DNA amplification from single cells. PCR-free library construction from the amplified gDNA proceeded following manufacturer’s protocols. A total of about 120 cells from tumor tissue and 4 cells from normal tissue were used for construction of scWGS libraries. The bulk DNA-seq libraries (containing about 1000 cells) were prepared exactly following the scWGS library preparation protocol. We prepared bulk DNA-seq libraries with 2 tumor samples and 1 normal sample. The scWGS libraries and bulk DNA-seq libraries were sequenced on Illumina NovaSeq6000 system, setting with paired-end 150bp.

### Data and samples

We generated scRNA-seq data of 9 samples from 3 colorectal cancer patients. We also generated scWGS data of 187 cells from tumor tissue and 5 cells from normal tissue from patient CRC#1, as well as bulk DNA-seq data of 2 tumor samples and bulk DNA-seq data from 2 normal samples from patient CRC#1. By integration the scRNA-seq data from Lee et al. (Lee et al. 2020), we obtained scRNA-seq data of 36 samples from 12 patients. The samples include 12 tumor tissues, 12 normal tissues, 9 tumor border tissues and 3 para-cancerous tissues.

### scRNA-seq data pre-processing

Cell Ranger v3.1.0 (10x Genomics) was used to perform reads alignment (human reference genome GRCh38), barcode demultiplexing, transcripts assemblies and expression counting, similar to our previous studies (Zhou and Jin 2020; Qin et al. 2021; Wang et al. 2022). The raw count matrices outputted by Cell Ranger were further processed by Seurat v3.2.1 package (Stuart et al. 2019). We filtered out the low-quality cells using the following criteria: 1) cell with <500 genes; 2) cell with <1000 UMIs; 3) cell with >50% mitochondrial genes; 4) potential doublets. We further removed plasma & red blood cell marker genes potentially caused by soup pollution.

### scWGS data pre-processing

The reads of scWGS were mapped to human reference genome GRCh38 using BWA VN:0.7.17 (Li and Durbin 2010). CNVs in each cell were identified by CNVkit-0.9.6 (Talevich et al. 2016). In brief, the scWGS library with extremely duplicated probes were considered as low quality, thus cells will be filtered out if the average depth of the 10 deepest contigs is greater than 10^2.5^ fold the median depth of the genome. We obtained the normalized CNV after the probes (log2 ratio) in each cell were segmented and smoothed by winsorize and multipcf in copynumber v1.15.0 R package (Nilsen et al. 2012). SNVs were identified by Mutect2 in GATK 4.0 (BD 2020), with bulk DNA-seq data of normal tissue as control (Panel of Normal). SNVs that detected in <3 cells will be filtered out.

### Dimension reduction and unsupervised clustering of scRNA-seq

scRNA-seq data was normalized using NormalizeData function in Seurat (Stuart et al. 2019) with default parameter. High-variable genes in scRNA-seq data were selected by FindVariableFeatures function in Seurat. We selected 2,000 high-variable genes for dimension reduction and unsupervised clustering. To reduce the impact of soup pollution, plasma and red blood cell marker genes (plasma cell marker genes include *IGHA1*, *IGHA2*, *IGHG1*, *IGHG2*, *IGHG3*, *IGHG4*, *IGHD*, *IGHE*, *IGHM*, *IGLC1*, *IGLC2*, *IGLC3*, *IGLC4*, *IGLC5*, *IGLC6*, *IGLC7*, *IGKC* and *JCHAIN*, red blood cell marker genes include *HBA1*, *HBA2*, *HBB*, *HBD*, *HBE1*, *HBG1*, *HBG2*, *HBM*, *HBQ1* and *HBZ*) were used to perform regression to get an unbiased z-score expression matrix by ScaleData function. Principal component analysis (PCA) was conducted on the unbiased z-score matrix. The top 20 principal components (PCs) were used to conduct Uniform Manifold Approximation and Projection (UMAP). Based on the 20 PCs, we clustered the cells using FindNeighbors and FindClusters with default parameter.

### Annotation of cell clusters

Cell clusters identified based on scRNA-seq data were annotated by cell type specific markers: immune cell (*PTPRC*), T cell (*CD3D*, *CD3E*), macrophage (*CD68*, *CSF1R*), mast cell (*MS4A2*, *TPSAB1*), B cell (*CD79A*, *CD79B*), plasma cell (*IGHA1*, *IGHG1*), endothelial cell (*PECAM1*, *VWF*), stromal cell (*COL1A1*, *DCN*), epithelial cell (*EPCAM*), and cancer cell (*MMP7*, *CEACAM6*). Those marker genes were essentially consistent with parikh et al. (Parikh et al. 2019).

### Inference of CNVs to identify cancer cells using scRNA-seq data

Normalized expression matrix was used to infer CNV in each cell. In detail, the genes with mean normalized expression level >0.1 across all cells were sorted by their genomic position. The genes were grouped into bins by applying a sliding window of 51 genes with a 10-genes step in each chromosome, in which each bin was called a probe. The mean of all genes’ z-score of normalized expression level in a probe was called the inferred raw CNV level. The raw inferred CNV level of a probe minus the average raw inferred CNV level in reference cells is the inferred CNV level of a probe.

By using stromal cells and endothelial cells as reference, the inferred CNV level matrix of all epithelial-like cells was obtained from normalized expression matrix in each patient. Then we define each cell’s inferred CNV score as the sum of the squared inferred CNV level of each probe in each cell. By comparing the mean value of all cells’ inferred CNV scores between cell clusters, we separated malignant cells and normal epithelial cells in each patient. Infercnv v1.12 (Patel et al. 2014) from Board Institute was also used to display patients’ CNV level.

### Identification of tumor subclone using scRNA-seq

To measure cell similarity at CNV level, the inferred CNV level matrix was processed for chromosome arm weighted PCA and unsupervised clustering. We apply hclust function with ward.D2 method on the top 10 PCs of inferred CNV level to perform cell clustering, and use cutree function to classify cells into tumor subclones. We set the number of tumor subclones (k) from 6 to 2 and chose the largest k as the number of tumor subclones when each tumor subclone has unique CNV pattern in each patient.

### Comparison of CNVs inferred by scWGS and CNVs inferred by scRNA-seq

Because we have both scWGS data and scRNA-seq data from patient CRC#1, we can compare whether the CNVs inferred by scRNA-seq are consistent with that inferred by scWGS. We mapped the high-resolution CNV regions inferred by scWGS to CNV regions inferred by scRNA-seq data for integration analysis. The integrated CNV level is the average CNV level weighted by its original CNV region percent. In order to assign the cells from scWGS data to the tumor subclones inferred by scRNA-seq data, we calculated the Elucidate distance of the cell to the centroid of each tumor subclone based on CNV level. The cell was assigned to the tumor subclone with the shortest distance to the cell.

### Pseudotime inferences of tumor subclones

We used UMAP_CNV_ in slingshot (Street et al. 2018) to calculate pseudotime of tumor subclones in CRC1. We use reduceDimension with DDRtree method in monocle (Qiu et al. 2017) to calculate pseudotime and embedded trajectory.

### Identification of SNVs in scWGS and scRNA-seq

We identified the SNVs in each cell from scWGS data following aforementioned method (BD 2020) (χ^2^ test; BH adjusted p value <0.05). We identified total 1,403 SNVs on autosome and X chromosome. For scRNA-seq data, we extracted cancer cells and epithelial cells from raw bam files that output from Cell Ranger. We called the SNVs in scRNA-seq data following our previous studies (Fu et al. 2021; Qin et al. 2021). We finally obtained 150 SNVs after removing recorded RNA editing sites in database RADAR (Ramaswami and Li 2014) and DARNED (Kiran and Baranov 2010). If the frequency of alterative allele is ≥1%, or if there is one alterative allele when total number reads is ≤100, the cell will be marked as an alternative cell at a this SNV. A SNV detected in >10 cells and had at least 5% alternative cells will be regarded as a validated SNV. A SNV with significantly different alternative cell ratio between tumor subclones (Wilcoxon test) will be regarded as tumor subclone specific SNV.

### Circos plot of CNV regions

We defined a CNV region as a duplication region or a deletion region when the mean CNV level of this region across all cells is larger or less than 0.01 in a patient. All patients’ ploidy CNV regions are shown on a circos plot (Krzywinski et al. 2009).

### Entropy and inferred CNV score of each tumor subclone

We used LandSCENT (version 0.99.3) (Teschendorff and Enver 2017) to calculate each cell’s entropy. The entropy and inferred CNV score of each tumor subclone are calculated by averaging the entropies and inferred CNV scores of all the cells in the subclone, respectively.

### Coefficient of variation (cell-cell variation) of gene expression

The coefficient of variation of gene expression was calculated for genes that expressed in more than half cells in cancer cells, epithelial cells, and each epithelial cell subtype which contain more than 100 cells and described in reference (Parikh et al. 2019). Undiff, Colonocytes, CT_colonocytes, Goblet, EECs and BEST4_OTOP2 among epithelial cell subtypes were defined by their marker genes, others were defined by high expressed gene in that cluster. There are 1440 genes expressed in more than 50% of cancer cells and at least 1 normal epithelial cell subsets. Wilcoxon Rank-Sum Test (paired samples) was used to test whether the coefficient of variations between two cell types/subtypes were significantly different.

### Relationship between gene expression and its DNA copy number

The expression level of each gene in a cell was calculated by dividing the UMI count of this gene by the total UMI count in this cell. The relative expression level of each gene in cancer cells was normalized by its expression level in epithelial cells. In brief, we calculated the fold change of gene expression in cancer cells relative to epithelial cells and clamped it between 0.2 and 5 to avoid the strong influence of outliers. Then, we divided genes that expressed in ≥5% cancer cells or normal epithelial cells into noCNV, deletion and duplication regions in each patient, and calculated average relative expression level of those genes in each category.

Correlation coefficient between gene expression level and its CNV level in cancer cells were calculated. Variations of CNV level in each CNV region across all cells were calculated in each patient. CNV regions were classified into 2 groups according to whether the variation >0.001 or not. Pearson’s correlation coefficient between the expression level of a gene and the CNV level of the gene located region cross all cells was calculated.

### Calculation of intra-tumoral heterogeneity scores

The intra-tumoral heterogeneity scores were calculated following Wu et. al., (Wu et al. 2021a). To calculate the intra-tumoral heterogeneity scores based on CNV (HetCNV), cell-cell similarities were calculated by Pearson’s correlation coefficients between the inferred CNV scores in each patient. Interquartile range (IQR) of the distribution of cancer cell pairs’ similarities was HetCNV. The intra-tumoral heterogeneity scores based on gene expression (HetGEX) were calculated similar to HetGEX but using normalized gene expression matrix.

### Identification of DEGs between cell types or cell subpopulations

DEGs between cell types or cell subpopulations were identified by FindMarkers function in Seurat. We set logfc.threshold =log2(1.5) to identify the DEGs between cancer cells and epithelial cells, or DEGs between tumor associated endothelial cells (or stromal cells) and their normal counterparts. The subclone specific genes for each tumor subclone in each patient were identified using FindAllMarkers with parameter logfc.threshold = log2(1.5) and logfc.threshold = log2(1.8). Enrichment analysis of each gene set was conducted using Metascape (Zhou et al. 2019).

### Separating DEGs between tumor subclones into two populations

The distribution of correlation coefficient between gene expression level and CNV level revealed the existence of two populations of DEGs. DEGs among tumor subclones were separated into two population at the lowest point of the distribution (r=0.29). To keep the number of genes with low CNV-expression correlation similar to that with high CNV-expression correlation, we set logfc.threshold = log2(1.5) for the genes with high CNV-expression correlation and logfc.threshold = log2(1.8) for the genes with low CNV-expression correlation. GO term enrichment in molecular function by enrichGO in clusterprofile (Wu et al. 2021b).

### Analysis of cell-cell communication

Cell-cell communication signatures and communication strengths were inferred by using cellchat v1.4.0 (Jin et al. 2021). There’re 8 patients among all 12 patients used in cell-cell communication analysis, other 4 patients are filtered due to few cells in many cell types or cells concentrated in few cell types. There are 78 cytokine signaling related pathways and 73 adhesion signaling related pathways for cytokine signaling analysis and adhesion signaling analysis, respectively. The significantly changed pathways between two cell types were calculated by rankNet function. The communication distance of a pathway is calculated by rankSimilarity function between normal and tumor sample in each patient. Interaction weight between cell types is drawn by netVisual_circle function.

## Acknowledgements

This study was supported by National Key R&D Program of China (2021YFF1200900, 2021YFA0909300), National Natural Science Foundation of China (32170646), Guangdong Basic and Applied Basic Research Foundation to N. H. (2023A1515011908), Shenzhen Science and Technology Program (KQTD20180411143432337), Shenzhen Innovation Committee of Science and Technology (JCYJ20220818100401003, ZDSYS20200811144002008). We thank all members from the Jin lab for the helpful discussion. We acknowledge the assistance of Core Facilities of SUSTech. The computational work was supported by Center for Computational Science and Engineering at SUSTech.

## Author contributions

W.J. conceived the project. Q.Z., K.L, Y.L. collected the clinical samples and performed experiments. W.C carried out the computational analysis with help from X.W., J.S., R.F.. W.J., E.C., N.H supervised the project. W.C. and W.J. prepared the manuscript, with all authors’ contribution.

## Data availability statement

The sequence data have been deposited in the Genome Sequence Archive in BIG Data Center under accession number HRA004167 and HRA000086. Code used for analysis is available https://github.com/CaoWei-UM/Single-cell-CRC-pipeline-2023.

---

**Supplementary Figure 1.**
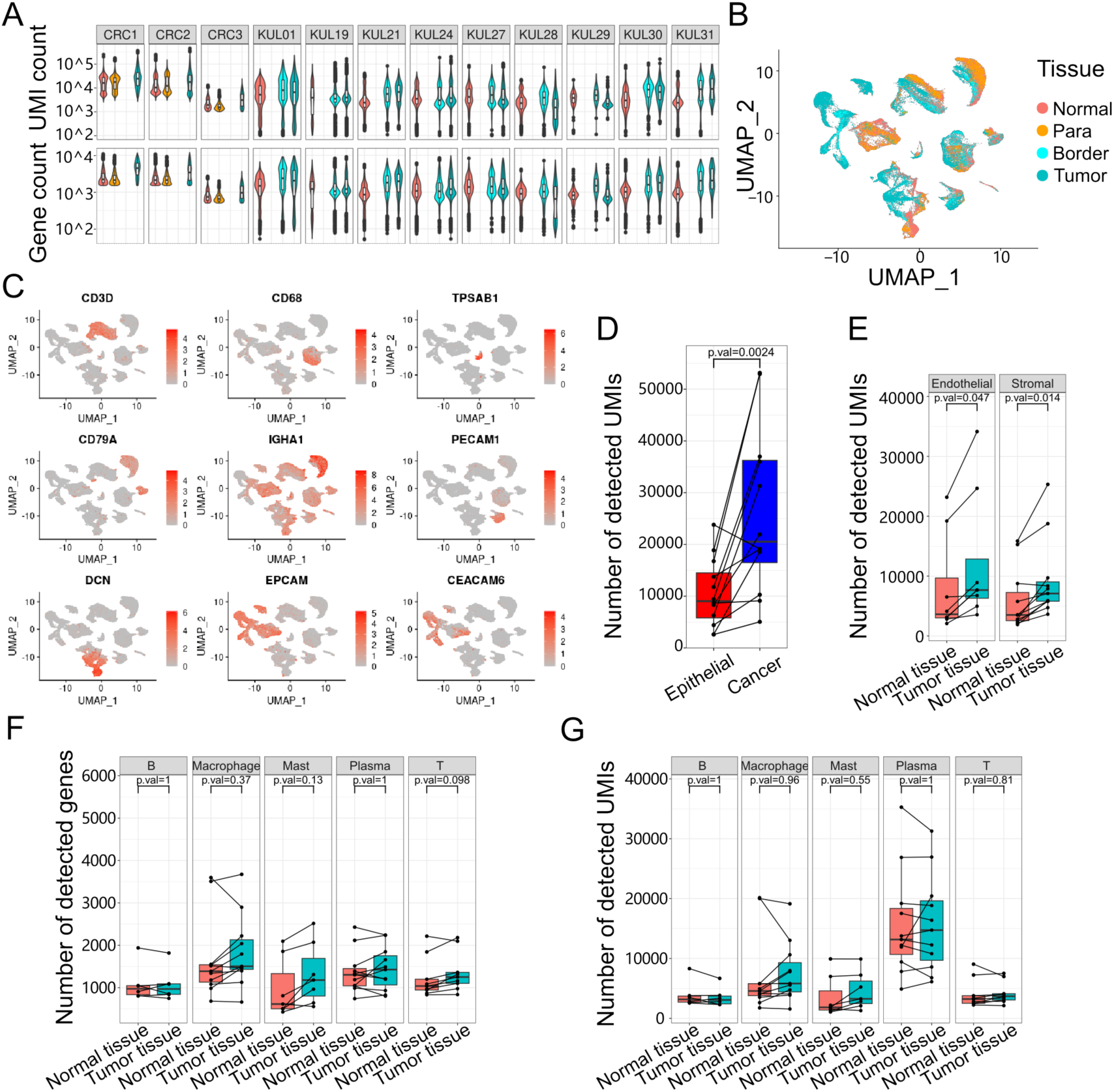
Sample, patient and cell types. **(A)** Violin plot of gene and UMI count of each sample in each patient. **(B)** UMAP visualization of 101,065 cells from tumor tissue and normal colorectal tissue in 12 patients, colored by sampling location. **(C)** UMAP visualization of markers’ expression levels for immune cells, stromal cells, epithelial cells, and cancer cells. **(D)** Boxplot of average number of detected UMIs in each patient for cancer cells and epithelial cells. Wilcoxon signed rank test were performed to examine whether the number of detected UMIs are significantly different between cancer cells and epithelial cells. **(E)** Boxplot of average number of detected UMIs in each patient in normal tissue and tumor tissue for endothelial cells and stromal cells. Wilcoxon signed rank test were performed to examine whether the number of detected UMIs were significantly different between normal tissue and tumor tissue for endothelial cells and stromal cells. **(F-G)** Box plot of average number of detected genes **(F)** and UMIs **(G)** and of each patient in immune cells for normal tissue (magenta) and tumor tissue (cyan). Paired Wilcoxon signed rank test was used for examining whether differences were significant.

**Supplementary Figure 2.**
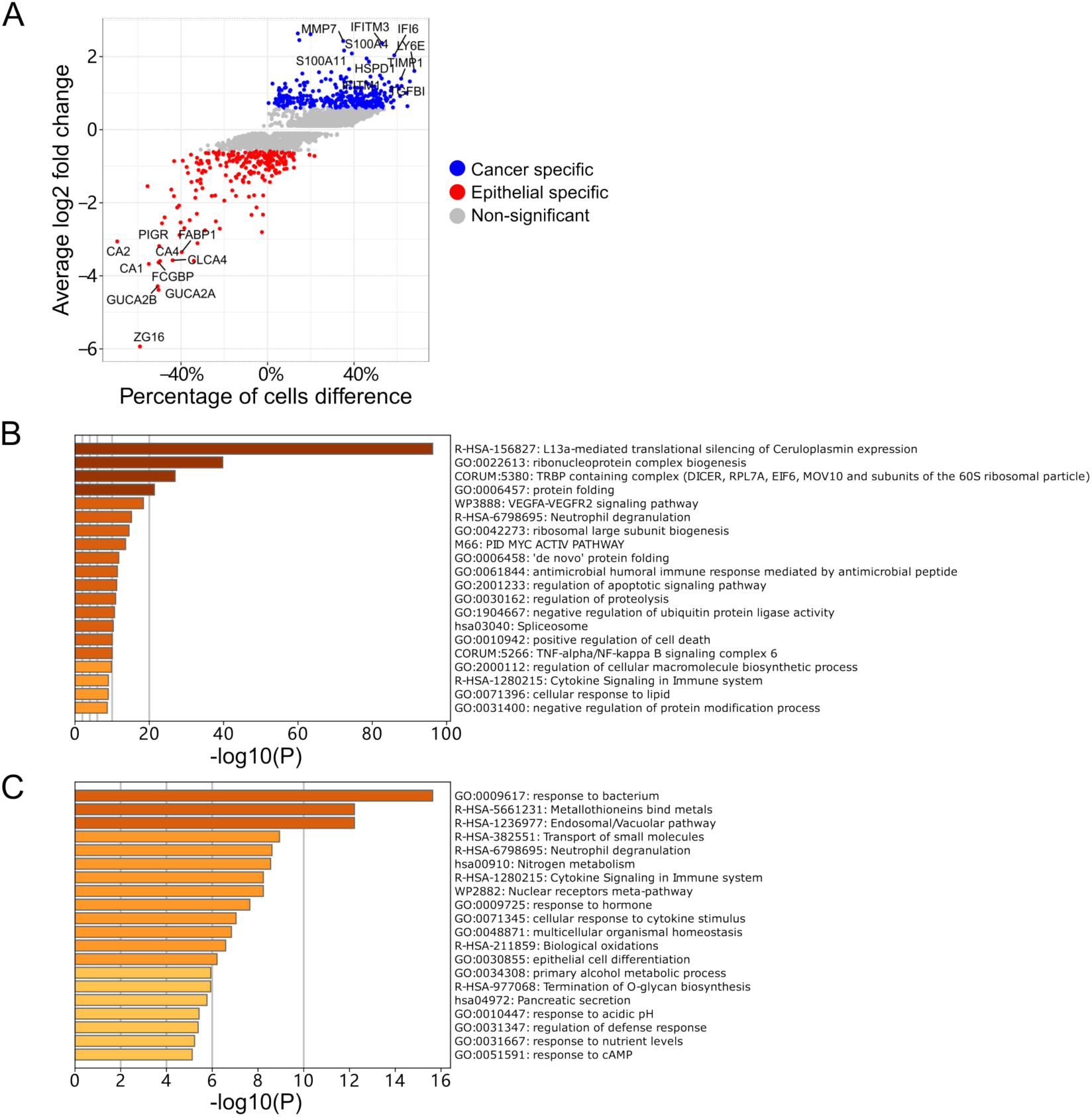
DEGs between cancer cells and epithelial cells. **(A)** Di-volcano plot of DEGs between cancer cell and epithelial cells in all patients. Blue and red represents cancer cell specific genes and epithelial cells specific genes. **(B)** Enrichment analysis of cancer cell specific genes. **(C)** Enrichment analysis of epithelial cells specific genes.

**Supplementary Figure 3.**
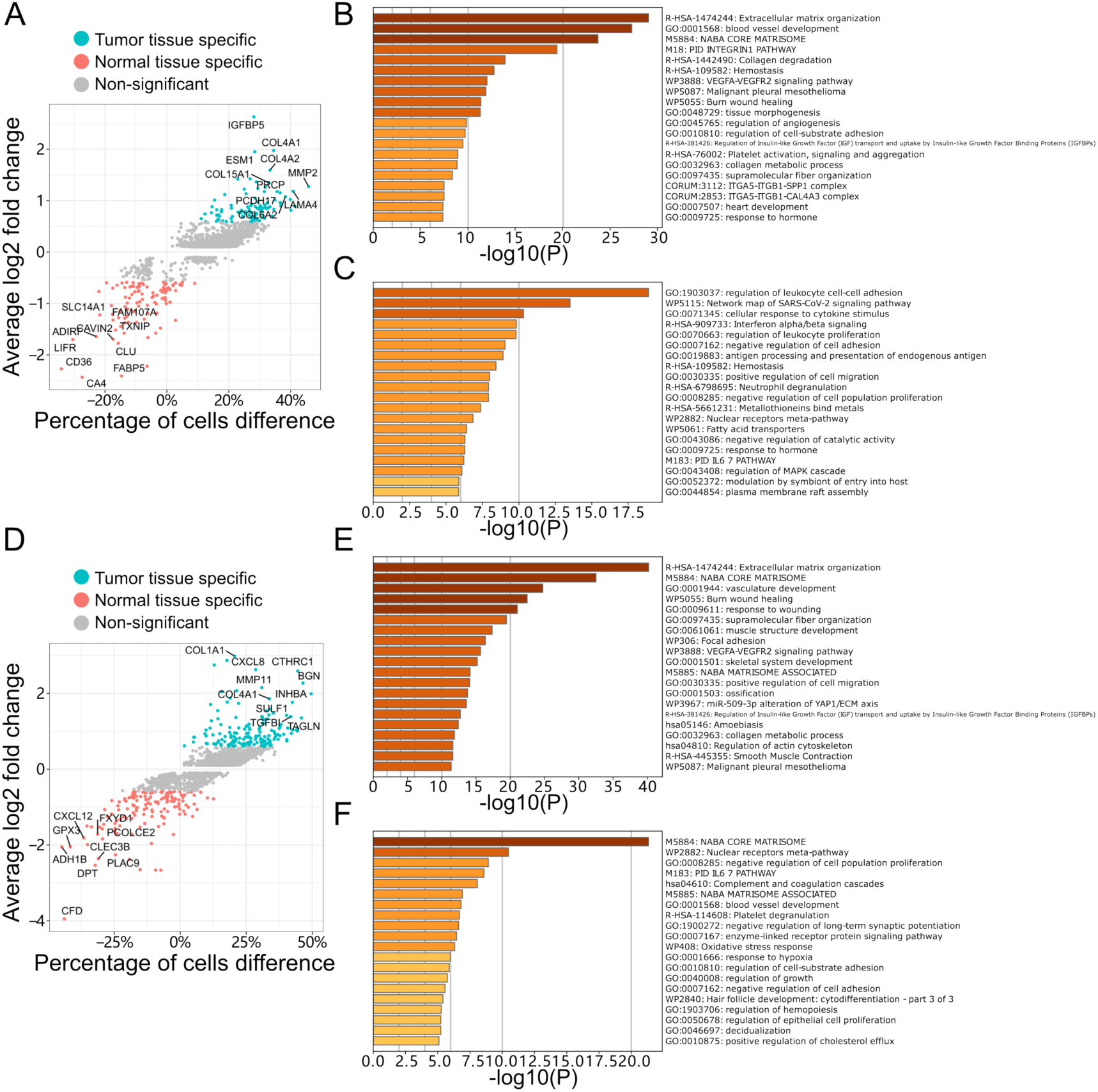
DEGs between tumor associated endothelial cells (or stromal cells) and their normal counterparts. **(A)** Di-volcano plot of DEGs between tumor associated endothelial cells and their normal counterparts. Cyan and magenta represent tumor associated endothelial cell specific genes and normal endothelial cell specific genes, respectively. **(B)** Enrichment analysis of tumor associated endothelial cells specific genes. **(C)** Enrichment analysis of normal endothelial cells specific genes. **(D)** Di-volcano plot of DEGs between tumor associated stromal cells and their normal counterparts. Cyan and magenta represent tumor associated stromal cell specific genes and normal stromal cell specific genes, respectively. **(E)** Enrichment analysis of tumor associated stromal cells specific genes. **(F)** Enrichment analysis of normal stromal cells specific genes.

**Supplementary Figure 4.**
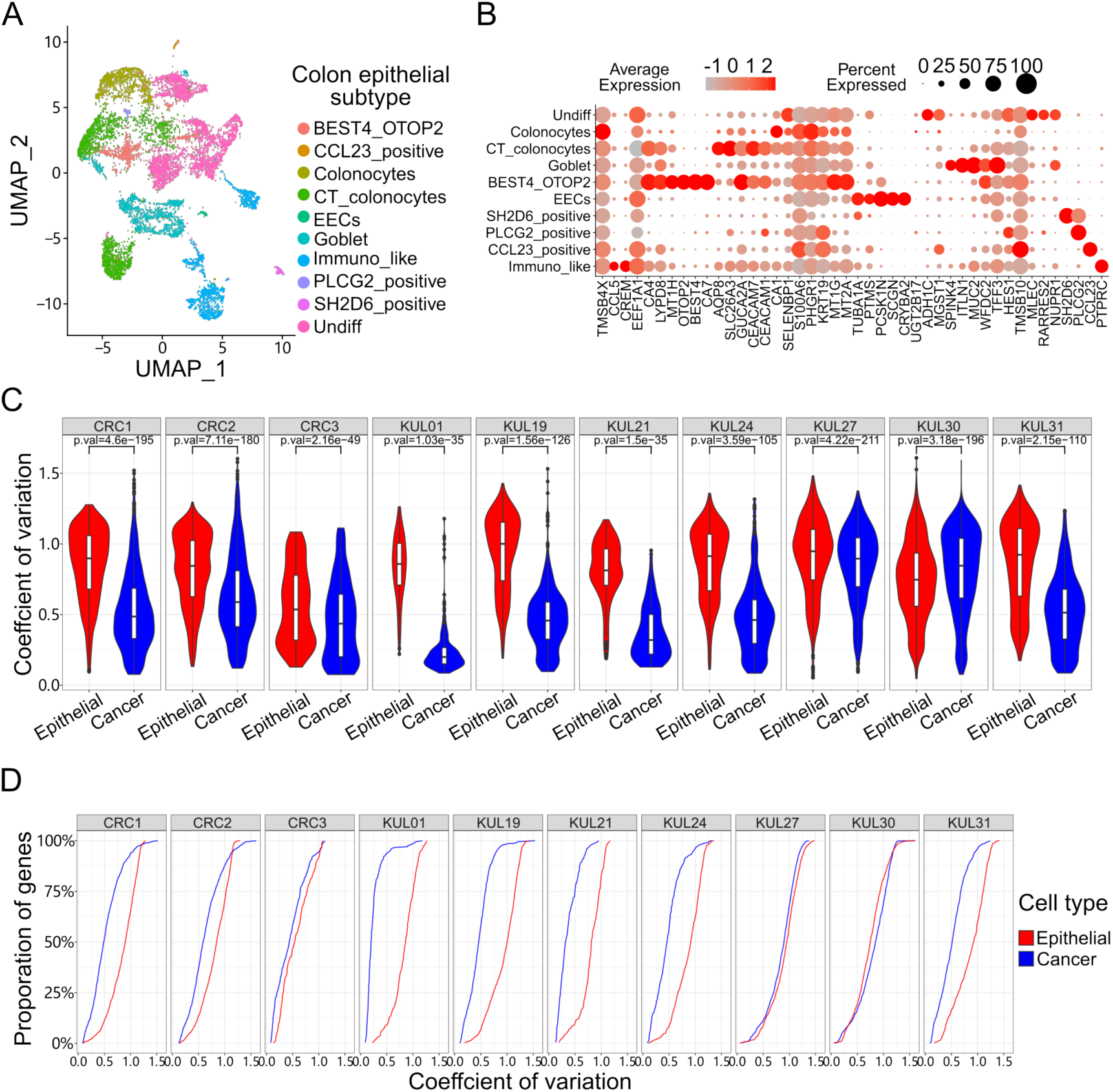
Epithelial cell subtypes and coefficient of variation of gene expression in epithelial cells. **(A)** UMAP visualization of 9,080 epithelial cells from normal colorectal tissue in 12 patients, colored by epithelial cell subtypes. **(B)** Dot plot of normalized expression level and expression percentage of epithelial cell subtype specific genes in 10 cell subtypes. **(C)** Violin plot of CV of gene expression in epithelial cells (red) and cancer cells (blue) in each patient. Paired Wilcoxon signed rank test was used to examine whether differences were significant. **(D)** Accumulation curve of CV of gene expression level in epithelial cells compared with same genes in cancer cells in each patient.

**Supplementary Figure 5.**
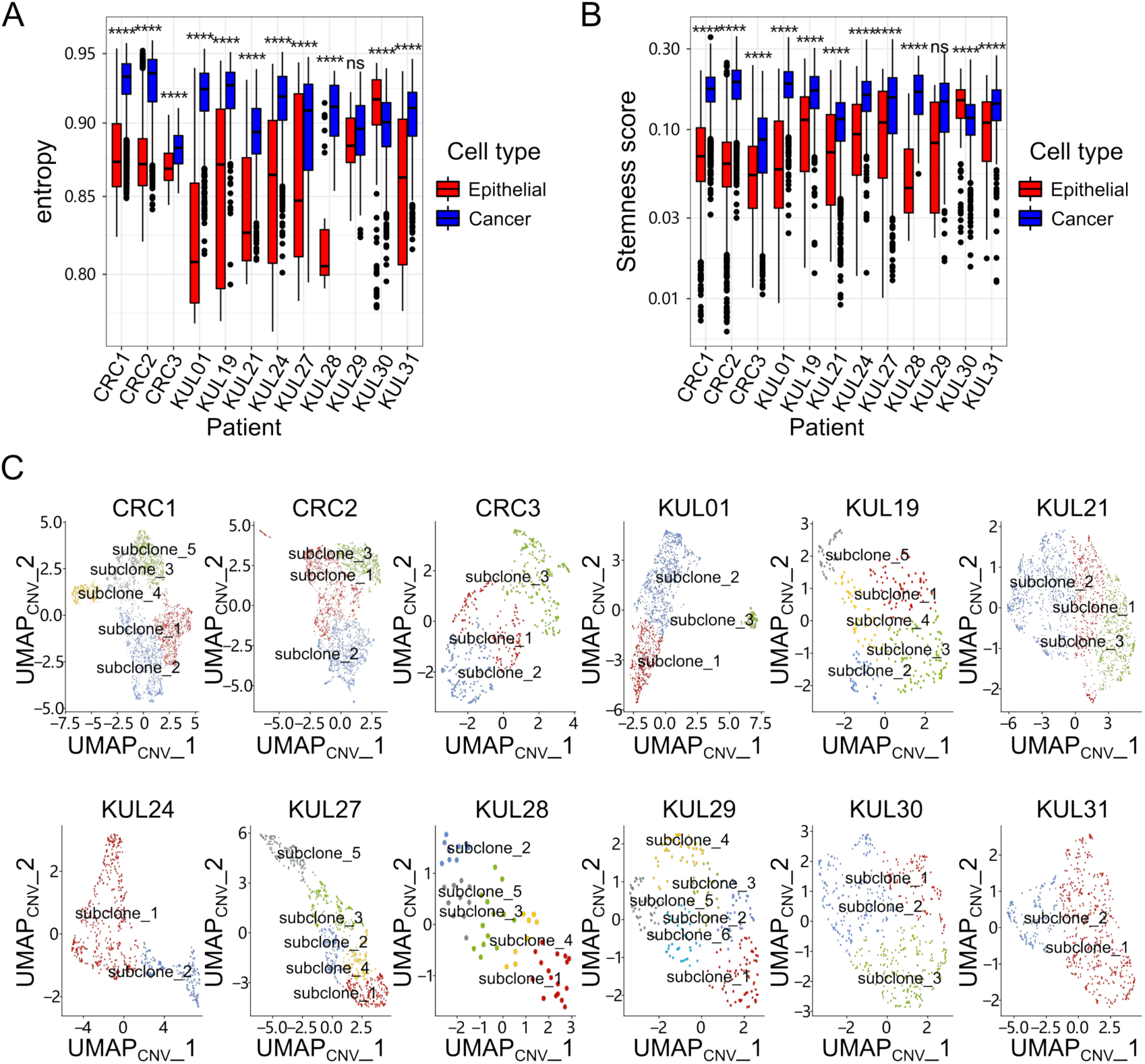
The heterogeneity of tumors and cancer cells. **(A)** Box plot of entropy in epithelial cells (red) and cancer cells (blue) in each patient. Paired Wilcoxon signed rank test was used for examining whether differences were significant. **(B)** Box plot of the stemness score in epithelial cells (red) and cancer cells (blue) in each patient. Paired Wilcoxon signed rank test was used for examining whether differences were significant. **(C)** UMAP visualization of cancer cells in each patient. The inferred CNVs in each cell was used for the dimension reduction and cell clustering. Cells were colored by tumor subclones.

**Supplementary Figure 6.**
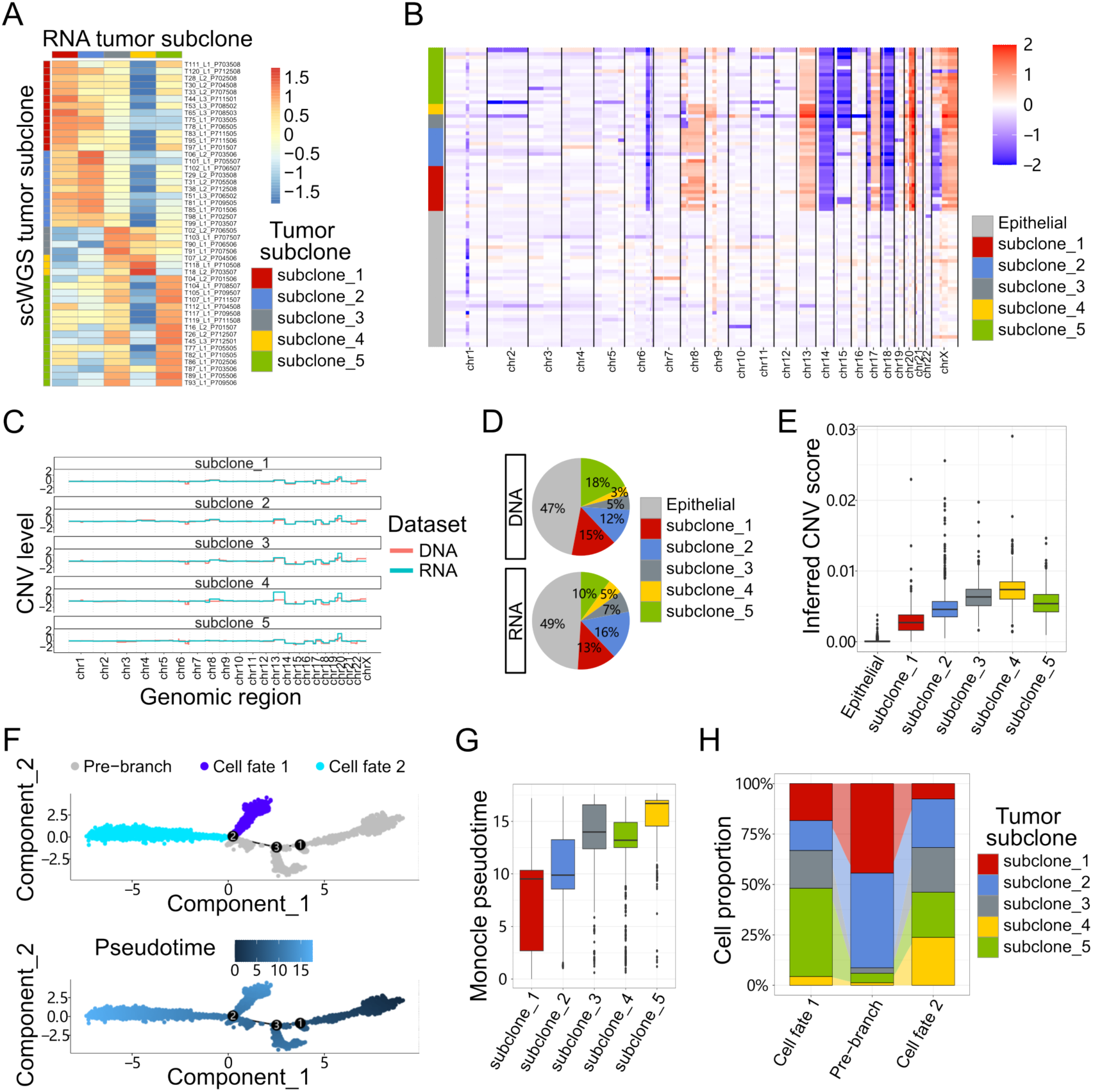
Integration of scWGS and scRNA-seq for analyses of tumor subclones and trajectory of CRC#1. **(A)** Heatmap shows tumor subclones inferred by scWGS data are similar to tumor subclones inferred by scRNA-seq in CRC#1. Cells are colored by similarities to tumor subclones. **(B)** Heatmap of CNV level in genomic regions in cells from scWGS data in patient CRC#1. Cells are grouped by epithelial cells and tumor subclones. **(C)** Comparison of CNV level between tumor subclones inferred by single cell RNA-seq data and their counterparts inferred by scWGS data. The CNV level was obtained by binning the CNV level into CNV regions across chromosomes. **(D)** Pie plot of proportion of non-cancer cells and tumor subclones in tumor tissue in scWGS data and scRNA-seq data. **(E)** Box plot of inferred CNV score in epithelial cells and each tumor subclone in patient CRC#1. **(F)** Lineage trajectories of cancer cells in CRC#1 inferred by monocle2, comparing with the lineage inferred by slingshot. **(G)** Box plot of pseudotime of cells in each tumor subclone in patient CRC#1 inferred by monocle2. **(H)** Percentage of tumor subclones in 2 lineage trajectories and the pre-branch trajectory.

**Supplementary Figure 7.**
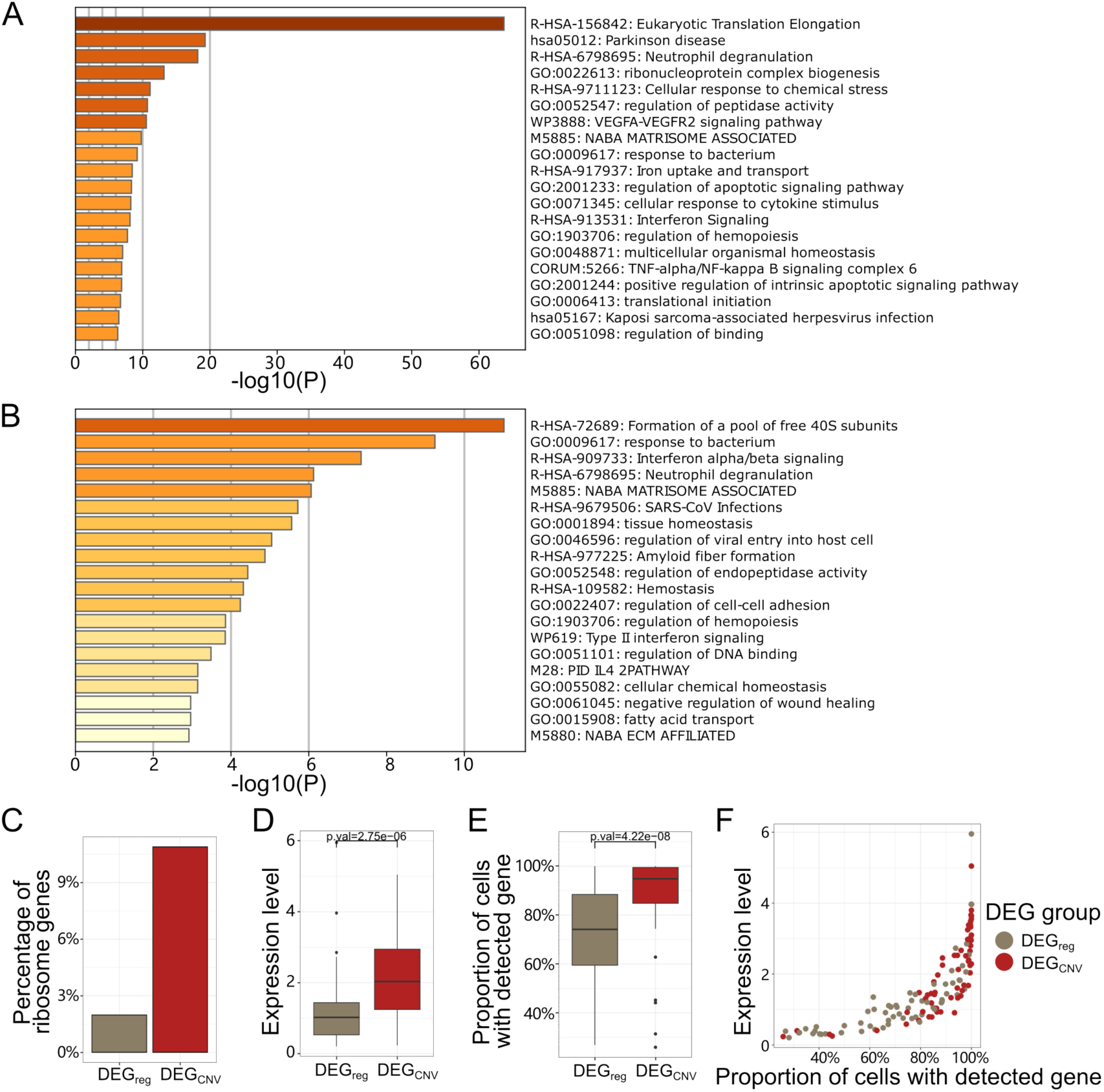
The differences of basic features between DEG_CNV_ and DEG_reg_. **(A)** Enrichment analysis of DEGs between tumor subclones (fold change > 1.5). **(B)** Enrichment analysis of DEGs between tumor subclones (fold change > 1.8). **(C)** Bar plot of the fraction of ribosome genes in DEG_CNV_ and DEG_reg_. **(D)** Box plot of expression level of DEG_CNV_ and DEG_reg_. Wilcoxon signed rank test was used to examine whether the differences were significant. **(E)** Box plot of proportion of cells with detected genes in DEG_CNV_ and DEG_reg_. Wilcoxon signed rank test was used to examine whether the differences were significant. **(F)** Scatter plot of proportion of cells with detected genes and expression level of DEG_CNV_ and DEG_reg_.

**Supplementary Figure 8.**
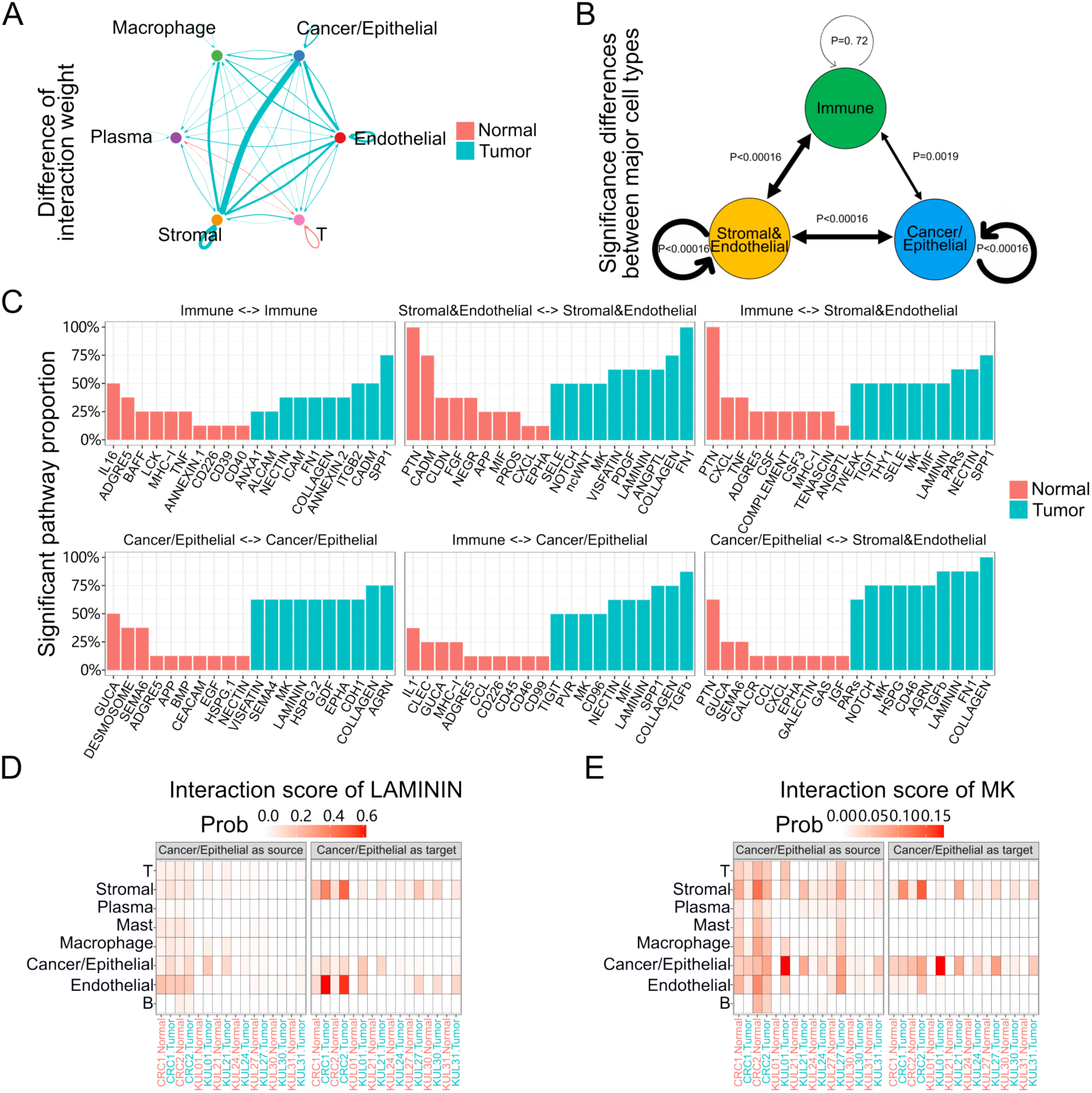
Cell-cell communication in tumor tissues and normal tissues. **(A)** Change of cell-cell communication (ligand/receptor pairs) network in tumor tissues comparing to normal tissues in all patients. Cyan and magenta represent increase and decrease in tumor tissue, respectively. **(B)** Diagram of change of cell-cell communication between immune cells (include T cell, plasma cell and macrophage), stromal&endothelial cells and cancer/epithelial cells. The interactions among any pair increases except the interaction between immune cells and immune cells. Paired Wilcoxon signed rank test were used to calculate the P value. **(C)** Bar plot of the top 10 pathways whose interaction weight significantly changed between normal tissues and tumor tissues in each major cell type pairs. The height of bars shows the proportion of the number of patients that involved in those pathways. **(D-E)** Heatmap of interaction score of cancer/epithelial cell to other cell types (left) and other cell types to cancer/epithelial cell (right) of LAMININ **(D)** and MK **(E)** pathways in each patient’s normal and tumor tissue.

